# Individual corticosterone response to intermittent swim stress predicts a shift in economic demand for ethanol from pre-stress to post-stress in male rats

**DOI:** 10.1101/2024.02.26.582066

**Authors:** Christopher L. Robison, Victoria Madore, Nicole Cova, Robert C. Drugan, Sergios Charntikov

## Abstract

This study investigated the relationship between stress exposure and subsequent ethanol use, focusing on individual differences among male rats. We combined operant self-administration with behavioral economics to assess how intermittent swim stress affects ethanol consumption. This approach allowed for a nuanced analysis of the transition from regular ethanol intake to stress-induced escalation in economic demand. Results showed a consistent rise in ethanol demand post-stress among subjects, irrespective of exposure to actual swim stress or a sham procedure. This increase may result from a two-week abstinence or an inherent rise in demand over time. Significantly, we identified a direct link between post-stress corticosterone levels and the demand for ethanol, considering baseline levels. This correlation was particularly pronounced when examining the shifts in both corticosterone levels and demand for ethanol post-stress. However, neither post-stress corticosterone levels nor their change over time correlated significantly with changes in ethanol demand following a forced swim test that was administered 24 hours after the intermittent swim stress test. This suggests potential context-specific or stressor-specific effects. Importantly, pre-stress ethanol demand did not significantly predict the corticosterone response to stress, indicating that high ethanol-demand rats do not inherently exhibit heightened stress sensitivity. Our research brings to light the complex interplay between stress and ethanol consumption, highlighting the critical role of individual differences in this relationship. This research introduces a nuanced perspective, underscoring the need for future studies in the realm of stress and substance use to give greater consideration to individual variability.

## 1. Introduction

Alcohol consumption ranks as the foremost cause of preventable deaths globally, accounting for approximately 3 million fatalities annually, or 5.3% of all deaths (World Health Organization, 2018). Beyond its health implications, alcohol use significantly contributes to social and economic burdens worldwide (Rehm et al., 2017; Thavorncharoensap et al., 2009). The link between traumatic stress events and alcohol use disorder is well-established, with evidence suggesting a strong correlation between stress and increased alcohol consumption (Keyes et al., 2012). However, existing preclinical studies, while affirming this link, often vary in outcomes based on factors like the animal model used, duration of alcohol access, and the characteristics of the test subjects. Traditional research models, such as group designs, provide valuable insights but may not fully capture the individual variability crucial in the development of stress-related alcohol use disorders. This gap highlights the need for more nuanced, individual-focused research to better understand the complex interplay between stress factors and alcohol use, particularly using clinically relevant paradigms.

Traumatic stress events significantly increase the risk of substance use, notably alcohol dependence, with high comorbidity observed in individuals with post-traumatic stress disorder (Breslau et al., 2003; Jacobsen et al., 2001; Mills et al., 2006). Such stress events not only contribute to the onset of alcohol use disorders but can also trigger relapse in those recovering from substance use. This necessitates a deeper investigation into the complex stress-alcohol use relationship (Dewart et al., 2006; Zywiak et al., 2003). Preclinical studies using animal models have been instrumental in exploring this association. Various methodologies, including free-choice home cage drinking and operant self-administration models, have provided insights into how different types of stress, like maternal separation, footshock, and social defeat, influence alcohol intake. These studies reveal that factors like the timing and intensity of stress exposure can have varying effects on alcohol consumption, underscoring the importance and role of individual variability in this context (Caldwell and Riccio, 2010; Caplan and Puglisi, 1986; Casey, 1960; Croft et al., 2005; Darnaudéry et al., 2007; Hilakivi-Clarke et al., 1991; Jaworski et al., 2005; Mills et al., 1977; Ploj et al., 2003; Roman et al., 2003; Siegmund et al., 2005; Van Erp and Miczek, 2001). Importantly, the variability in responses seen in these studies highlights the need for further research focusing on individual differences in stress and ethanol use interactions.

Recent studies have unveiled a common neurobiological pathway connecting stress responses in the brain to the efficacy of anxiolytic drugs. The Learned Helplessness (LH) model suggests that stress enhances the brain’s response to drugs targeting the benzodiazepine/GABA receptor complex. This is supported by a range of research, including some of our work, showing stress-related alterations in this receptor complex across various species and individual subjects (Braestrup et al., 1979; Drugan et al., 2016, 1996, 1989a, 1989b, 1985, 1984; Havoundjian et al., 1986; Maier and Seligman, 1976). These findings also show the variability in pharmacological responses to stress. Our study’s methodology, based on the intermittent swim stress model and reinforcer demand modeling, is sensitive to these pharmacological effects and aims to capture the individual differences in behavioral and neurobiological responses to stress (Brown et al., 2001; Christianson and Drugan, 2005; Drugan et al., 1992; R. C. Drugan et al., 2013; Robert C. Drugan et al., 2013; Stafford et al., 2019; Warner et al., 2013a, 2013b). The reinforcement demand modeling is grounded in microeconomic theory and has been applied to study behavioral responses to various stimuli (Hursh et al., 2005; Hursh and Roma, 2016). By training rats to respond for a reinforcer under varying fixed ratio (FR) schedules, we can assess their willingness to work for different reinforcers (Killeen and Jacobs, 2017; Schwartz et al., 2021). This methodology, which has been refined in prior work from our laboratory, can be used to offer insights into individual substance use patterns (Kazan et al., 2020; Robison et al., 2023; Stafford et al., 2019, 2019). The present study’s phased approach, involving initial chronic ethanol self-administration followed by stress exposure and reassessment of economic demand for ethanol, offers a distinctive and pertinent perspective on how stress affects alcohol consumption at an individual level.

The primary goal of this study is to explore how intermittent swim stress interacts with ethanol self-administration at an individual level. We hypothesized that increased stress sensitivity, indicated by elevated corticosterone levels relative to baseline, would correlate with a higher economic demand for ethanol post-stress. To test our hypothesis, we designed a study in which rats initially engaged in ethanol self-administration and then were subjected to intermittent swim stress, followed by an assessment of their stress-induced ethanol demand. This design diverges from typical preclinical models that introduce stress before ethanol exposure, more accurately reflecting the sequence of events in human experiences with trauma and alcohol use. The integration of the intermittent swim stress and reinforcer demand modeling offers a robust framework for correlating individual stress responses with ethanol demand shifts, enhancing our understanding of stress-alcohol interaction and informing future individual-level research.

## 2. Materials

### 2.1. Subjects

Forty-two male Wistar rats weighing between 250-300 g were obtained from Envigo (Indianapolis, IN, USA). All rats were included in the study. The rats were individually housed in a temperature-controlled vivarium with a 12-hour light/dark cycle, with lights turning on at 0700. After being introduced to the colony, the rats were allowed to acclimate for one week before the commencement of the experimental procedures.

During this acclimation period, the rats had access to food and water ad libitum. Throughout the rest of the study, the rats were subjected to food restriction to maintain their weight at 90% of their free-feeding weight, with water available without restriction. The free-feeding weight was gradually increased by 2 g every 30 days. All procedures were conducted in accordance with the Guide for the Care and Use of Laboratory Animals (National Research Council et al., 2010) and were reviewed and approved by the University of New Hampshire Institutional Animal Care and Use Committee.

### 2.2. Apparatus

#### 2.2.1. Self-administration chambers

Behavioral tests were conducted in sound- and light-attenuated Med Associates conditioning chambers (30.5 × 24.1 × 21.0 cm; l × w × h) equipped with an exhaust fan (ENV-018MD; Med Associates, Inc.; St. Albans, VT, USA). The chambers had aluminum sidewalls, metal rod floors, and polycarbonate surfaces. Two retractable levers (147 nN required for micro-switch closure) were mounted on each side of the right-side wall and were used as manipulanda to operate the retractable sipper (ENV- 252M; Med Associates, Inc.; St. Albans, VT, USA) positioned on the wall between those levers. Cue lights were positioned above each lever. Med Associates interface and software (Med-PC for Windows, version IV) were used to collect data and execute programmed events.

#### 2.2.2. Intermittent Swim Stress

Intermittent swim stress was conducted in two acrylic cylinders (21 cm × 42 cm; d × h) with a 6.35 mm galvanized wire mesh at the bottom of each cylinder. Cylinders were suspended over a tank (80.6 cm × 45.7 × 28.6 cm; l × w × h) filled with water maintained at 15 ± 1◦C. The apparatus was controlled by a Med-PC interface and software (Med Associates Inc., St. Albans, VT, USA). Space heaters, above and in front of each cylinder, circulated warm air (∼36◦C) in and around the cylinders to limit the effects of hypothermia during the inter-trial intervals.

#### 2.2.3. Forced Swim Test

The forced swim test was conducted in acrylic cylinders (20 cm × 46 cm; d × h). The water was filled to 30 cm height and was kept at 24°C.

### 2.3. Drugs

Ethanol (200 proof; Decon Labs; King of Prussia, PA, USA) and sucrose (store-bought sugar) solutions were made using tap water.

## 3. Methods

Figure 1 presents the experimental progression.

**Figure 1.**
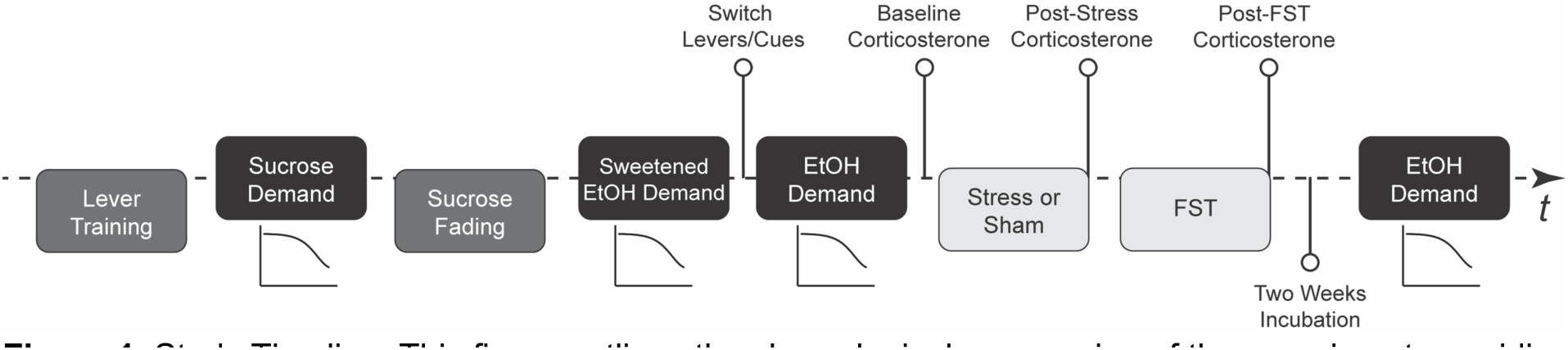
Study Timeline. This figure outlines the chronological progression of the experiment, providing the main stages of the research procedure (N=42; 32 rats in stress condition and 10 rats in sham stress condition). FST stands for forced swim test, EtOH for ethanol, and *t* is for time.

### 3.1. Lever Training

The rats were initially trained to consume a 12% (w/v) sucrose solution from a retractable sipper during sessions that lasted 120 minutes. These sessions involved non-contingent delivery of sucrose rewards on a variable time interval, approximately three rewards per minute. Next, the rats were trained to press a lever to obtain the 12% sucrose solution through an auto-shaping procedure. Each session commenced with the illumination of cue lights, the extinguishing of the house light, and the insertion of a randomly selected lever (either right or left). Pressing the lever or a lapse of 15 seconds resulted in the following consequences: the sipper tube was inserted, the lever was retracted, the cue lights located above each lever were turned off, and the house light was turned on. After 5 seconds, the sipper tube was retracted, the cue lights were re-illuminated, the house light was extinguished, and a randomly selected lever was reinserted into the chamber. The program ensured that the same lever was not presented more than two times consecutively, and the number of presentations of the left and right levers was kept equal throughout the session. Training persisted until the rats pressed the levers on at least 80% of lever insertions for two successive days. This resulted in total training times ranging between three to six daily sessions, depending on individual performance.

### 3.2. Acquisition of economic demand for 12 % sucrose

Rats were pseudo-randomly assigned active levers, ensuring an even split between right and left. They were trained over three consecutive days to self-administer 12% sucrose on a fixed reinforcement schedule (FR1). Each session started with cue lights on, house light off, and the insertion of levers. Upon meeting the active lever’s schedule requirement, the following occurred: the sipper tube was inserted, levers were retracted, cue lights turned off, and the house light was turned on. After 5 seconds, this sequence was reversed, and the levers were reinserted. These self-administration sessions, lasting 4 hours, took place during the day cycle (0900-1500). Following three days of 12% sucrose self-administration, the rats were put on a daily escalating fixed ratio (FR) reinforcement schedule, using the sequence: 1, 3, 5, 8, 12, 18, 26, 38, 58, 86, 130, 195, 292, 438, and 657. This continued until they failed to earn at least one reinforcer.

Finally, the rats self-administered 12% sucrose on a variable reinforcement schedule (VR3; range 1-5) until all rats completed the demand assessment, followed by three additional sessions to reestablish 12% sucrose self-administration.

### 3.3. Sucrose fading

Rats were trained to self-administer ethanol via a sucrose-fading procedure across 4- hour sessions, keeping active lever assignments from the previous phase. The protocol mirrored the previous phase with adjusted liquid reinforcers. Initially, rats received a 12% sucrose solution, gradually fortified with increasing ethanol concentrations every four days in the sequence: 2%, 4%, 8%, and 12%. After six consecutive days with 12% sucrose and 12% ethanol, sucrose was progressively reduced to 2% over stages (12%, 8%, 4%, and 2%, each for four days). Following the fading protocol, rats self-administered a mixture of 2% sucrose and 12% ethanol (referred to later as sweetened ethanol) using a VR3 reinforcement schedule.

### 3.4. Acquisition of economic demand for sweetened ethanol (2 % sucrose and 12 % ethanol solution)

The acquisition of economic demand for sweetened ethanol followed the same procedure as that for 12% sucrose, but with 2% sucrose and 12% ethanol solution as the reinforcer. Afterward, rats underwent self-administration of 2% sucrose and 12% ethanol solution on a VR3 schedule of reinforcement until all rats completed demand assessment and an additional three daily sessions to reacquire self-administration behavior.

### 3.5. Acquisition of economic demand for ethanol-alone

The 2% sucrose and 12% ethanol solution was replaced with a 12% ethanol-alone solution. Active lever assignments were switched, and associated cues were changed to eliminate any potential sucrose reward-related conditioning. Each ethanol-only session began with the house light on and levers inserted. Upon meeting the schedule requirement, the sipper tube was inserted, levers retracted, the house light was turned off, and cue lights were turned on. After five seconds, this sequence was reversed, allowing for continued lever pressing for ethanol. Following this protocol, rats self-administered ethanol-alone on a VR3 reinforcement schedule for 7 to 10 daily 4-hour sessions until active lever presses outnumbered inactive ones. Economic demand acquisition for ethanol-alone followed the previously described 12% sucrose protocol. Once the terminal schedule requirement was reached, where rats failed to earn at least one reinforcer, all rats were allowed to self-administer ethanol-alone on a VR3 schedule of reinforcement until all rats completed demand assessment plus an additional three daily sessions to reacquire ethanol self-administration. All sessions took place during the day cycle.

### 3.6. Exposure to a stress episode: intermittent swim stress

Each swim trial consisted of a 5-s forced swim in which the cylinder was submerged to a depth of 25 cm. Eighty trials were presented at a variable 60 s (10–110 s) inter-trial-interval. Immediately following intermittent swim stress, rats in the stress condition were towel-dried and returned to the vivarium to heated cages (heating pad placed underneath cages). Rats in the sham stress condition underwent the same procedures, with the exception that the apparatus was not filled with water and rats were not towel-dried. This sham stress condition served as a control to compare against the stress exposure, enabling us to assess the impact of stress on corticosterone response.

### 3.7. Examining the carry-over effects of prior stress exposure: forced swim test

The forced swim test was administered 24 hours following the intermittent swim stress exposure. During the forced swim test, all rats were forced to swim for 5 min in 24 ± 1◦C. Water depth was kept at 30 cm. This depth level forced rats to swim because it avoids the possibility of body support by the tail touching the bottom of the cylinder (i.e., tail-standing). The forced swim test was included 24 hours after the intermittent swim stress as it serves a key purpose in examining the carry-over effects of prior stress exposure on subsequent stress responses. By implementing the forced swim test 24 hours after the intermittent swim stress, we can determine whether prior stress exposure intensifies, dampens, or has no effect on the corticosterone response to a subsequent stressor on an individual level and then correlate these effects with economic demand for ethanol. Furthermore, the forced swim test is a standardized stressor that elicits a reliable hormonal response, making it a suitable choice for this type of analysis.

### 3.8. Blood collection and analysis

Blood was collected from rats before stress exposure (baseline), 30 min after intermittent swim stress, and 30 min post forced swim test, aligning with peak corticosterone concentrations (Connor et al., 1997; Stafford et al., 2019). Blood ethanol concentration tests occurred immediately after one hour of ethanol-alone self-administration that substituted a regular 4-hour session. There were two plasma alcohol concentration tests separated by at least two days of ethanol-alone self-administration. Rats were gently wrapped in a towel, and their tails were immersed in 46 ± 2◦C water to facilitate vasodilation. Using a #11 scalpel, a small incision was made in the tail, and about 300 microliters of blood was gathered via a capillary tube, all within a 3-min window. Following blood collection, rats were returned to their home cages.

Subsequent incisions were at least 1 cm away from previous ones (Drugan et al., 2005; Stafford et al., 2019). Samples were centrifuged at 4◦C for 4 min at 1,300 rpm, and plasma was separated and stored at -80°C for later assay. Enzyme-linked immunoabsorbent assays (Arbor Assays, Ann Arbor, MI, USA) were used to analyze samples in duplicates, with corticosterone concentrations read at 405 nm using a BioTek microplate reader with Gen5 software. An Ethanol Assay Kit (ab65343; Abcam; Cambridge, UK; McCarter et al., 2017) was used to measure the average plasma alcohol concentration from both samples for each rat (the average was used for analyses).

### 3.9. Data analysis

The economic demand for a reinforcer was assessed using the Exponential Model proposed by Hursh and Silberberg (Hursh and Silberberg, 2008). Consumption data (g/kg) from each reinforcement schedule were fit into the nonlinear least squares regression model using the formula: 𝑙𝑜𝑔𝑄 = 𝑙𝑜𝑔𝑄0 + 𝑘 ×, 𝑒^(−αQ0 *C*)^ − 1). Here, *Q* denotes quantity consumed, *Q_0_* indicates quantity consumed when the price is zero (i.e., consumption at zero cost or maximal consumption), 𝑘 is a parameter that adjusts the range of the dependent variable (*logQ*), *e* is the base of the natural logarithm, *C* is the cost, and α is the rate of decline in consumption as cost increases (demand elasticity; smaller values indicate higher demand). The model estimated the demand elasticity (*α*) and intensity (*Q_0_*). Maximum expenditure (*O_max_*) was calculated using the highest expenditure for each price or reinforcement schedule. The point of price where demand becomes elastic, and expenditure reaches maximum (*O_max_*) is represented by *P_max_*. The Essential Value (*EV*), calculated as 1/(100 × α × k ^1.5^), inversely proportional to *α*, was derived from the economic demand model. *EV* quantifies a reinforcer’s ability to maintain operant behavior amidst escalating behavioral costs and is often used to signify the intensity of demand or the value of a commodity. The economic demand curve analysis was performed using GraphPad Prism version 9, specifically employing a Prism template created by Hursh, S. R., & Roma, P. G. (2014) for exponential demand curve modeling, which was downloaded from https://ibrinc.org/software/. This approach incorporates the use of an extra sum-of-squares F test, as recommended by the model, to compare the fit of curves via the alpha (α) parameter, providing a preliminary evaluation of general effects. Adopting this method aligns our analysis with a rigorously validated framework, ensuring consistency with established research practices. For a more detailed investigation, subsequent analyses, including ANOVAs and regression analyses that explore variations in essential values among other variables, were conducted. These comprehensive statistical evaluations, along with post-hoc multiple comparisons tests, were also carried out using GraphPad Prism software.

In our study, we use the Essential Value (*EV*) as a primary measure that captures key elements of the demand equation. These include consumption at different reinforcer prices and demand curve elasticity. While the behavioral economics model can yield a multitude of measures for investigating diverse aspects of behavior, our current study is not intended for such a broad exploration. The study was not designed nor initially planned for an exhaustive analysis of all potential measures derived from the behavioral economics model; thus, the unnecessary examination could introduce type 1 errors.

With that said, our primary interest in this study was to gauge each rat’s individual motivation for ethanol, as represented by the *EV,* and then compare it to other stress-related variables. We’ve successfully applied this approach in previous studies to evaluate both grouped and individual demand and then compare these results with other measures (Kazan et al., 2020; Kazan and Charntikov, 2019; Stafford et al., 2019). Therefore, in this study, we employ the *EV* as the main measure of economic demand for ethanol.

## 4. Results

### 4.1. Blood ethanol concentration

Our analysis confirmed a significant positive relationship between the volume of ethanol consumed and blood ethanol concentration (β = 0.006969, 95% CI: 0.006067 to 0.007871, p < 0.0001; R² = 0.7424), validating the reliability of this measure for consumption.

### 4.2. Descriptive statistics of economic demand data

Table 1 displays descriptive statistics by Reinforcer and Group, including mean, standard deviation (SD), and standard error of the mean (SEM) for *EV*, *α*, *O_max_*, and *P_max_*. These metrics are provided to offer a comprehensive overview of the data, although not all are directly utilized in subsequent analyses for the reasons described in the Data Analysis section.

**Table 1:** Descriptive Statistics of Behavioral Economic Parameters by Reinforcer and Group

| Reinforcer | Group | Mean (SD; SEM) |  |  |  |
| --- | --- | --- | --- | --- | --- |
| | | *EV* | $\alpha$ | *O<sub>max</sub>* | *P<sub>max</sub>* |
| Sucrose | Sham-Stress | 6.09 (2.84; 0.90) | 6.30e-05 (2.81e-05; 8.90e-06) | 314.34 (146.57; 46.35) | 23.56 (14.51; 4.59) |
| Sucrose | Stress | 7.71 (3.39; 0.60) | 5.07e-05 (2.51e-05; 4.44e-06) | 398.23 (175.29; 30.99) | 31.13 (19.61; 3.47) |
| Sweetened-EtOH | Sham-Stress | 0.52 (0.27; 0.08) | 1.49e-03 (2.75e-03; 8.70e-04) | 26.88 (13.69; 4.33) | 15.30 (5.10; 1.61) |
| Sweetened-EtOH | Stress | 0.56 (0.44; 0.08) | 1.25e-03 (1.80e-03; 3.19e-04) | 28.69 (22.63; 4.00) | 19.42 (15.92; 2.81) |
| EtOH-Pre-Stress | Sham-Stress | 0.12 (0.14; 0.04) | 0.02 (0.04; 0.01) | 6.43 (7.01; 2.22) | 8.84 (6.32; 2.00) |
| EtOH-Pre-Stress | Stress | 0.11 (0.12; 0.02) | 0.02 (0.04; 6.34e-03) | 5.92 (6.38; 1.13) | 7.38 (3.61; 0.64) |
| EtOH-Post-Stress | Sham-Stress | 0.20 (0.26; 0.08) | 0.03 (0.06; 0.02) | 10.46 (13.59; 4.30) | 10.51 (6.20; 1.96) |
| EtOH-Post-Stress | Stress | 0.19 (0.19; 0.03) | 0.02 (0.05; 9.25e-03) | 9.73 (9.78; 1.73) | 16.65 (37.78; 6.68) |

Note. EV = Essential Value; SD = Standard Deviation; SEM = Standard Error of the Mean.

### 4.3. Comparing demand for sucrose, sweetened ethanol, and ethanol-alone

First, we assessed demand curves for sucrose, sweetened ethanol, and ethanol-alone prior to stress, employing the extra sum-of-squares F test. Significant differences in the elasticity parameter alpha were detected across the three conditions [F(2,34) = 284, p < 0.0001; Figure 2A]. Specifically, the demand for sucrose was found to be the least elastic (alpha = 4.3e-005; recall that smaller values indicate higher demand), while the demand for ethanol prior to stress tests was the most elastic (alpha = 0.0030). The sweetened ethanol exhibited intermediate elasticity (alpha = 0.00057).

**Figure 2:**
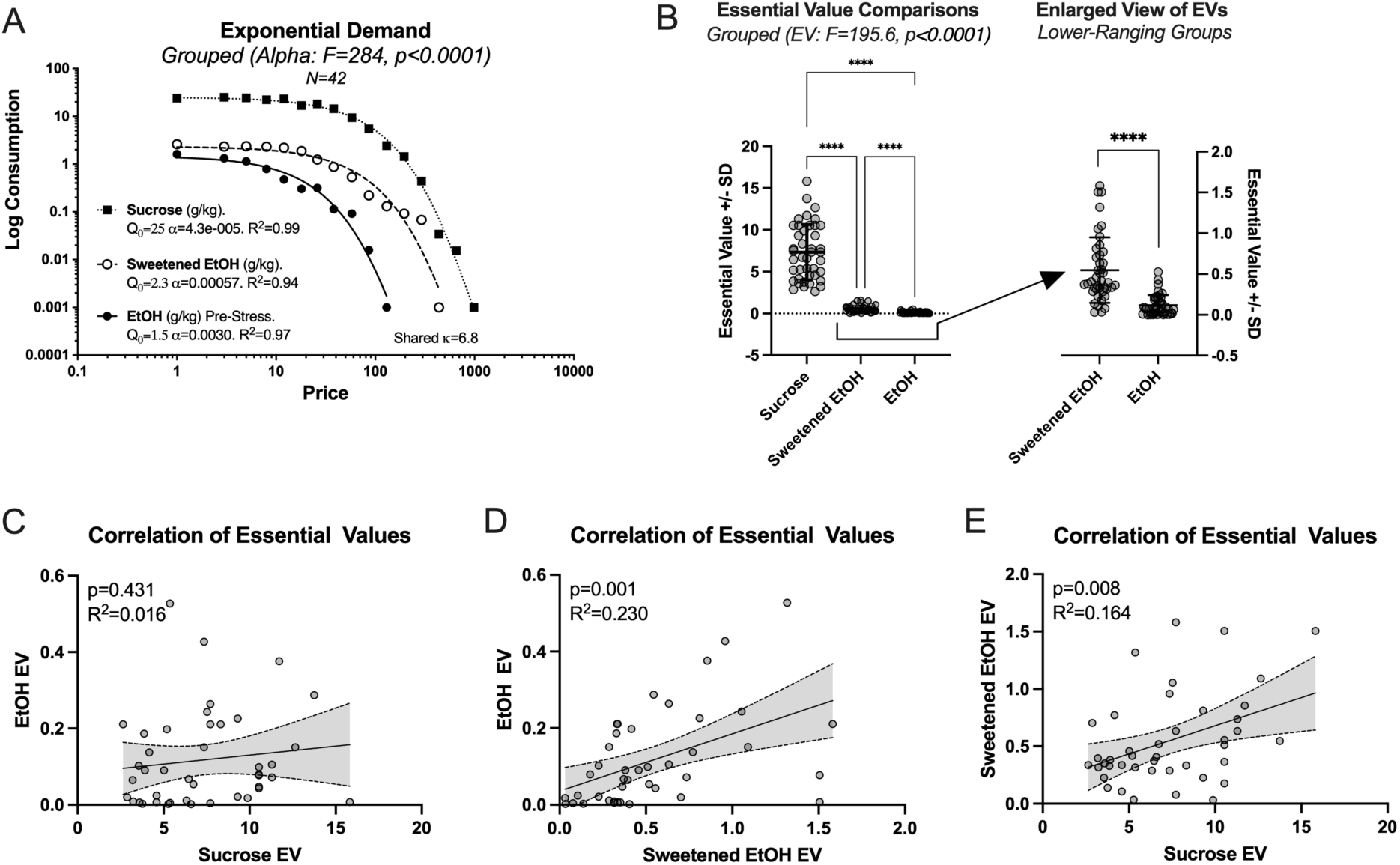
This figure presents a detailed analysis of the variations in demand elasticity and essential values (EV) across three conditions: sucrose, sweetened ethanol (EtOH), and ethanol-alone (N=42; within-subjects comparisons). (A) Represents significant differences in the elasticity parameter alpha across the conditions, with sucrose exhibiting the highest elasticity and ethanol-alone the lowest. (B) Demonstrates significant differences in the essential values across the three conditions, with all pairs showing significant mean differences (**** p<0.0001). (C) Shows no correlation between the essential values for responding to sucrose and ethanol-alone on an individual level. (D) Depicts a significant correlation between the essential values for responding to sweetened ethanol and ethanol-alone. (E) Highlights a significant correlation between the essential values for responding to sucrose and sweetened ethanol.

Following the observation of a significant main effect of elasticity, we conducted more refined analyses focusing on essential values. We employed a repeated measures one-way ANOVA with Greenhouse-Geisser correction, which revealed significant variations in essential values across three conditions [F(1.016, 41.66) = 195.6, p < 0.0001], accounting for approximately 82.67% of the variation in the response (R² = 0.8267; Figure 2B). Tukey’s post-hoc comparisons revealed significant mean differences between all treatment pairs: sucrose vs. sweetened ethanol (Mean Diff. = 6.777, p < 0.0001), sucrose vs. ethanol-alone (Mean Diff. = 7.207, p < 0.0001), and sweetened ethanol vs. ethanol (Mean Diff. = 0.4301, p < 0.0001).

On an individual level, the essential values for responding to sucrose and ethanol-alone were not significantly correlated. Simple linear regression showed a non-significant slope (β = 0.004706, 95% CI: -0.007238 to 0.01665, p = 0.4306) and a low coefficient of determination (R² = 0.0156; Figure 2C).

A significant correlation was observed between the essential values for responding to sweetened ethanol and ethanol-alone. Simple linear regression showed a significant slope (β = 0.1492, 95% CI: 0.06200 to 0.2365, p = 0.0013), and the model accounted for 23.01% of the variability in the response (R² = 0.2301; Figure 2D).

Moreover, a significant correlation was identified between the essential values for responding to sucrose and sweetened ethanol. This relationship was characterized by a significant slope (β = 0.04905, 95% CI: 0.01367 to 0.08442, p = 0.0078), with the model explaining 16.41% of the variability in the response (R² = 0.1641; Figure 2E).

### 4.4. Assessing the effects of stress on economic demand for ethanol

To assess the impact of stress on the economic demand for ethanol, changes in the alpha parameter (elasticity) were analyzed before and after exposure to cold swim stress, exclusively in rats within the stress condition, utilizing the extra sum-of-squares F test. Results showed a trend toward variability in elasticity, though it fell short of statistical significance [F(1,16) = 3.7, p = 0.0738, Figure 3A], suggesting no conclusive evidence to support differences in alpha across conditions. A comparative analysis of line fits for rats exposed to sham stress showed no significant change in the elasticity of ethanol demand (alpha) before and after the sham stress episode [F(1,17) = 1.5, p = 0.2330; Figure 3B].

**Figure 3:**
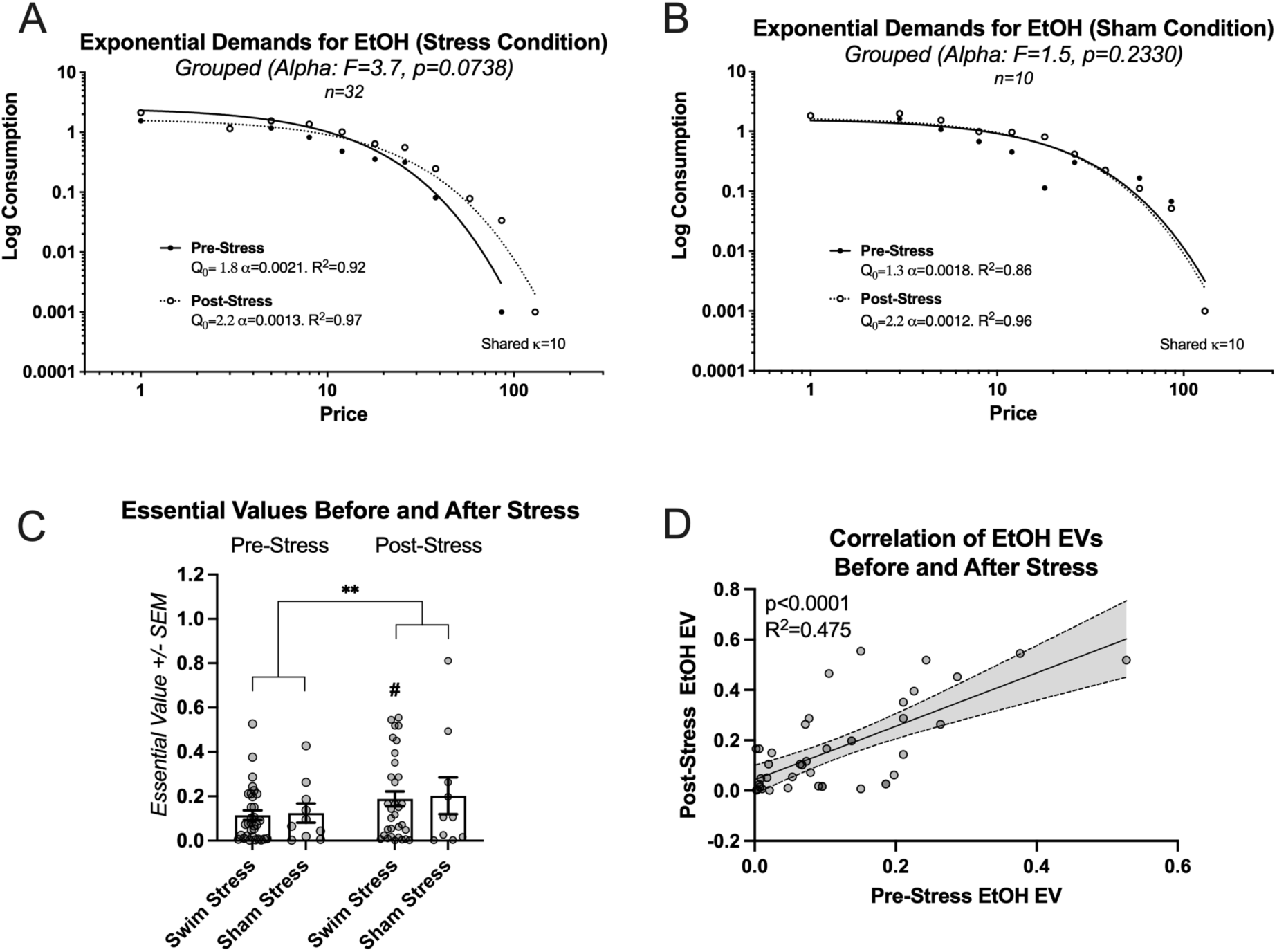
This figure details the changes in ethanol (EtOH) demand elasticity and essential value (EV) before and after the stress episode. (A) Illustrates the trend towards change in ethanol demand elasticity in rats subjected to intermittent swim stress, albeit not statistically significant. (B) Shows no significant alteration in ethanol demand elasticity in rats exposed to sham stress. (C) Illustrates that, although no significant interaction was found between the type of stress condition (swim stress vs. sham stress) and the testing times (pre- vs. post-stress), there was a statistically significant increase in the level of effort all rats dedicated to obtaining ethanol after experiencing stress. This is indicated by the main effect of time on ethanol-seeking behavior (p<0.01). # indicates significantly higher EV post-stress in comparison to pre-stress. (D) Shows a significant positive relationship between individual EVs before and after the stress episode for all rats (N=42).

We then conducted a more refined analysis focusing on the essential values. Two-way repeated measures ANOVA of essential values before and after intermittent swim stress episode demonstrated no significant interaction between the factors group (swim stress and sham stress) and time [pre-stress and post-stress; F(1, 40) = 0.006356, p = 0.9369; η² = 0.000027], and no significant main effect of group [F(1, 40) = 0.04484, p = 0.8334; η² = 0.0009]. A significant main effect of time was observed [F(1, 40) = 8.242, p = 0.0065; η² = 0.0355; Figure 3C]. Rats that were exposed to intermittent swim stress worked harder for ethanol after the stress exposure (EV; p=0.0134; Šídák’s multiple comparisons test; see # sign indicating this effect in Figure 3C). Because there was no effect of group, we proceeded with a correlation of individual essential values before and after stress episode for all rats.

A linear regression analysis of individual essential values before and after stress episode for all rats revealed a significant positive relationship, with a slope (β) of 1.060 (95% CI: 0.7045 to 1.416, p < 0.0001). This model accounted for 47.54% of the variability in the dependent variable (R² = 0.4754; Figure 3D).

### 4.5. Grouped corticosterone effects

To examine the impact of stress on a physiological benchmark of stress, such as corticosterone levels, we adopted a systematic and controlled comparison. This involved comparing effects across different groups (those exposed to stress and those not) and at various times (from baseline, following a stress episode, to after a forced swim test). Using a two-way analysis of variance (ANOVA), we identified significant main effects for both the stress condition [swim stress vs. sham stress; F(1, 120) = 8.434, p = 0.0044; η² = 0.048] and time [baseline, stress episode, forced swim test; F(2, 120) = 15.74, p < 0.0001; η² = 0.179], as well as a significant interaction between stress condition and time [F(2, 120) = 3.921, p = 0.0224; η² = 0.044].

Tukey’s post-hoc comparisons revealed a significant rise in corticosterone levels in rats exposed to swim stress and subsequently to the forced swim test compared to baseline (see Swim Stress bars in Figure 4; all p values are shown in the figure). However, for rats in the sham stress condition, their corticosterone levels did not significantly differ from baseline following sham stress. After the forced swim test, rats in the sham-stress condition showed significantly elevated corticosterone levels compared to baseline and post-sham stress time points (refer to Sham Stress bars in Figure 4). Notably, the corticosterone response in the swim stress group was significantly greater than that in the sham stress group following the stress or sham stress exposure (refer to the middle pair of bars in Figure 4).

**Figure 4:**
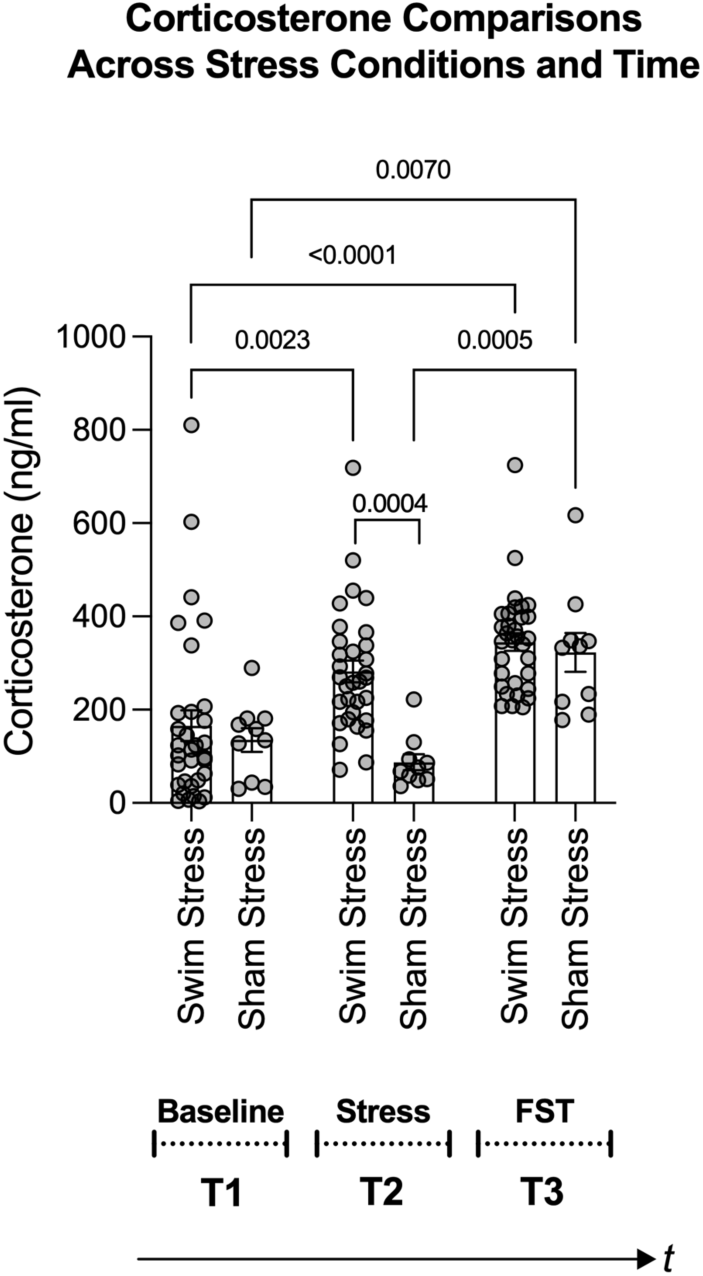
Figure shows changes in corticosterone levels based on stress condition (swim stress vs. sham stress) and time points: T1 (baseline), T2 (post swim/sham stress), and T3 (post forced swim test; FST). Notably, rats exposed to swim stress (n=32) showed a significant rise in corticosterone levels after the swim stress and FST. Rats in the sham stress condition (n=10) only showed a significant increase after the FST. The middle pair of bars underscores a greater corticosterone response in the swim stress group versus the sham group. Brackets with p-values atop indicate statistically significant differences.

### 4.6. Using individual corticosterone response to stress to predict escalation of demand for ethanol

After evaluating the effects of stress on corticosterone levels across groups, we shifted our focus to individual-level analyses. Our objective was to determine if the physiological stress response of an individual could predict an increase in the demand for ethanol. This aspect forms the core of our study, representing the primary analyses conducted throughout the research.

Corticosterone levels after intermittent swim stress did not significantly correlate with the post-stress essential value for ethanol, as indicated by a non-significant slope (β = 7.853e-005, 95% CI: -0.0004520 to 0.0006090, p = 0.7645) and a small coefficient of determination (R² = 0.003037; Figure 5A). This finding aligns with our initial prediction, as individual variability in baseline corticosterone levels among rats can mask the actual impact of stress on physiology. We posited that by focusing on the change in corticosterone response, calculated by subtracting baseline levels from post-stress levels, we could more accurately capture the individual’s physiological response to stress. This “delta” measure effectively normalizes each rat’s corticosterone response, reducing the confounding effect of baseline variability and providing a clearer picture of the stress response at an individual level.

**Figure 5:**
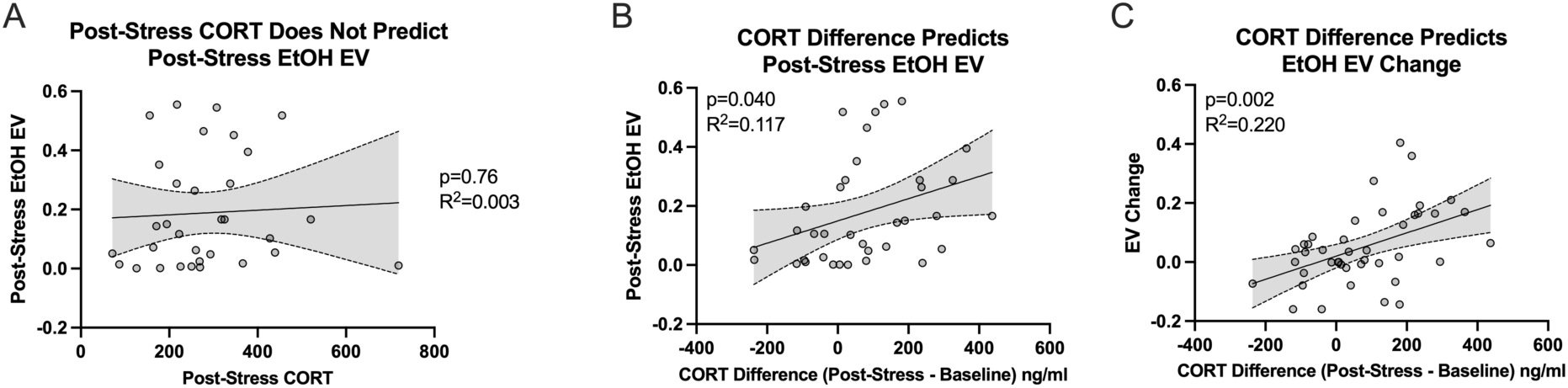
Figure shows the relationship between individual corticosterone (CORT) response to stress and the escalation in demand for ethanol (EtOH). (A) Shows no significant correlation between corticosterone levels post-swim stress and the post-stress essential value (*EV*) for ethanol (n=32). (B) Demonstrates a significant correlation when considering the change in corticosterone levels (post-stress minus baseline) with the post-stress essential value for ethanol (n=32). (C) Highlights a significant association between the change in corticosterone levels and the shift in essential value for ethanol (post-stress EV minus pre-stress EV; n=32).

In line with our hypothesis, the difference in corticosterone levels (post-intermittent swim stress minus baseline) showed a significant correlation with the post-stress essential value for ethanol. This correlation was denoted by a significant slope (β = 0.0003765, 95% CI: 1.764e-005 to 0.0007353, p = 0.0403) and a modest coefficient of determination (R² = 0.1179; Figure 5B). Furthermore, the change in corticosterone levels (post-intermittent swim stress minus baseline) had a significant association with the shift in essential value for ethanol (post-stress *EV* minus pre-stress *EV*). This relationship was signified by a significant slope (β = 0.0003935, 95% CI: 0.0001573 to 0.0006298, p = 0.0017) and a moderately high coefficient of determination (R² = 0.2208; Figure 5C). Importantly, using the shift in the essential value for pre-stress to post-stress doubled the amount of variance explained by the change in corticosterone response from before to after the stress episode. These findings underscore the importance of examining shifts or escalations in ethanol demand following stress exposure, reinforcing the pertinence of our focus on difference scores. Additionally, considering the dearth of studies exploring the effect of stress on subsequent ethanol intake, our research offers novel insight and highlights the necessity of further inquiry into how stress events may enhance the drive to consume ethanol.

Finally, we found no significant correlation between either the corticosterone levels after the forced swim test or the change in these levels (from baseline to post-test) and the demand for ethanol following the stress exposure or the change in ethanol demand from before to after the stress exposure. Specifically, there was a no significant correlation between post-forced swim test corticosterone and post-stress essential value for ethanol (β = 3.915e-005, 95% CI: -0.0006201 to 0.0006984, p = 0.9043; R²=0.0004900). Similarly, the correlation between the corticosterone difference and post-stress ethanol demand change was also not significant (β = 0.0001803, 95% CI: - 2.118e-005 to 0.0003818, p = 0.0780; R²= 0.07560).

### 4.7. Economic demand for ethanol prior to stress episode does not relate to corticosterone response to stress

No significant correlation was observed between the pre-stress essential value for ethanol and the corticosterone after the intermittent swim stress. The slope of this relationship was not statistically significant (β = 162.4, 95% CI: -234.0 to 558.9, p = 0.4094), and the coefficient of determination was low (R² = 0.02281; data not shown). Similarly, the post-forced swim test corticosterone levels did not significantly correlate with the pre-stress essential value for ethanol, as demonstrated by a non-significant slope (β = 31.14, 95% CI: -291.8 to 354.1, p = 0.8452) and a very low coefficient of determination (R² = 0.001291; data not shown). The change in corticosterone levels from baseline to post-intermittent swim stress also did not significantly correlate with the pre-stress essential value for ethanol, with a non-significant slope (β = 94.11, 95% CI: - 361.9 to 550.1, p = 0.6764) and a very low coefficient of determination (R² = 0.005886; data not shown). The absence of a significant correlation between corticosterone levels and pre-stress ethanol demand underscores that rats with high ethanol demand are not inherently more stress-sensitive. This distinction suggests that the observed post-stress escalation in ethanol demand may stem from stress exposure itself rather than from pre-existing individual differences.

## 5. Discussion

Research on alcohol consumption in rodents indicates that it is influenced by genetic, developmental, and social factors (Anacker and Ryabinin, 2010; Crabbe et al., 2006; Weinberg et al., 2008). Discrepancies in findings on stress and ethanol consumption likely stem from variations in research methodologies, including differences in ethanol models, stressors, duration of stress exposure, and assessment techniques (for reviews, see Logrip et al., 2012; Noori et al., 2014). The lack of research on how stress affects ethanol consumption at the individual level highlights the need for a spectrum of research strategies to address this gap. Our study contributes to this area by employing a behavioral economics framework, a 4-hour daily operant ethanol self-administration model, and cold-water swim stress. This approach arranges research phases in a clinically relevant manner: chronic ethanol self-administration, stress exposure, and post-stress ethanol demand evaluation. Our findings indicate no significant change in ethanol demand elasticity at the group level post-stress (Figure 3A). However, all rats, irrespective of stress condition, showed increased ethanol demand post-stress (Figure 3D), possibly due to an extended two-week abstinence period or an inherent increase in ethanol demand over time. Swim stress evoked higher corticosterone levels compared to sham stress (Figure 4 middle bars), validating our stress induction model. Our data reveal that post-stress corticosterone levels were not directly correlated with post-stress essential value for ethanol (Figure 5A). However, when controlling for baseline corticosterone variability, a significant relationship emerged between post-stress corticosterone changes and ethanol essential value (Figure 5B, C). These findings highlight the complexities in the relationship between stress and ethanol-taking behavior and underscore the importance of individual-level analysis in understanding these phenomena.

In the early phase of our study, we employed the sucrose fading protocol to facilitate operant ethanol self-administration. This protocol enabled us to gather key data providing insights into the interaction between individual primary reward preferences and ethanol preference. Results showed that rats worked hardest for sucrose, followed by sweetened ethanol and ethanol-alone (Figure 2B). Our findings deviate from previous research by showing higher elasticity in sucrose demand compared to sweetened ethanol and ethanol-alone (Figure 2A), suggesting differences in reinforcement properties (Heyman, 2000, p. 200, 1997, 1993; Heyman et al., 1999; Kim and Kearns, 2019; Petry and Heyman, 1995; Samson and Lindberg, 1984). This discrepancy might be due to experimental design differences or the unique characteristics of our subjects. Notably, our study also explores individual variability in demand for these reinforcers, a perspective rarely examined in previous research, with only one study from our own laboratory utilizing a similar approach but with a long-access ethanol self-administration model (Robison et al., 2023). We observed distinct patterns of effort exertion for sucrose, sweetened ethanol, and ethanol-alone at an individual level. The absence of a correlation between sucrose and ethanol-alone consumption indicates a significant qualitative difference in the transition to ethanol-alone consumption (Berridge, 2004; Kampov-Polevoy et al., 1999). Thus, our results reveal new dimensions to our understanding of the relationships between these substances not evident in group-level analyses (Bickel et al., 2012).

Our study further examined the effect of stress on ethanol demand by comparing two groups: one exposed to swim stress and a control group subjected to sham stress.

Despite a substantial sample size, detecting statistical effects in the stress-exposed group was challenging, suggesting limitations in conventional group-based approaches. Following stress exposure, both groups demonstrated heightened efforts to obtain ethanol. This increase could be attributed to the enforced abstinence period or a gradual escalation in demand for ethanol during the access phase, phenomena previously documented in other studies (O’Dell et al., 2004; Siegmund et al., 2005; Vengeliene et al., 2003). A strong correlation was observed between individual essential values before and after stress, indicating persistent ethanol preference despite stress interventions (Lesscher et al., 2010). These findings suggest that individual-level analysis may be suitable for a deeper understanding of the nuanced effects of stress on ethanol consumption.

The main objective of our study was to explore the impact of swim stress on individual ethanol demand. We used corticosterone levels as a biomarker of stress, reflecting activation of the HPA axis (Commons et al., 2017; Nishimura et al., 1988; O’Connor et al., 2003; Stafford et al., 2019). Our findings demonstrate a significant elevation in corticosterone levels following swim stress (Figure 4). Importantly, when accounting for baseline corticosterone variability, the post-stress corticosterone response predicted changes in ethanol demand (Figure 5C). Rats experiencing more stress, indicated by higher corticosterone levels post-swimming, exhibited increased efforts to obtain ethanol. This finding highlights the importance of selecting relevant variables in such research. Notably, direct comparisons between post-stress corticosterone and ethanol demand were less insightful, with baseline variations potentially masking stress effects. For instance, the correlation between post-stress corticosterone changes and ethanol-seeking was minor (11% variance explained). However, when comparing changes in corticosterone levels from baseline to post-stress with changes in ethanol demand from pre- to post-stress, the explanatory power increased to 22%. This suggests that a more nuanced analysis of stress biomarkers in relation to substance use behavior can yield deeper insights into the complex mechanisms underlying substance use.

Our findings indicate that individual responses to stress play a crucial role in alcohol use, underlining the importance of observing behavioral patterns at a personal level. This perspective could inform preventative measures, suggesting that addressing potential substance use issues before they arise post-stress may offer a more effective strategy than current practices. By tailoring interventions to individual experiences of stress, our research supports a proactive approach in the management of substance use and stress-related disorders.

## Acknowledgments

This work and S. Charntikov received partial support from NIGMS (Grant GM113131) and a NIDA/NIGMS joint grant (DA056871).

